# Chromosomally Encoded *mcr-5* in Colistin Non-susceptible *Pseudomonas aeruginosa*

**DOI:** 10.1101/297028

**Authors:** Erik Snesrud, Rosslyn Maybank, Yoon I. Kwak, Anthony R. Jones, Mary K. Hinkle, Patrick Mc Gann

**Affiliations:** Multi-drug resistant organism Repository and Surveillance Network, Walter Reed Army Institute of Research, Silver Spring, MD.

**Author notes:** Address correspondence to Patrick Mc Gann.

## Abstract

Whole genome sequencing (WGS) of historical *Pseudomonas aeruginosa* clinical isolates identified a chromosomal copy of *mcr-5* within a Tn*3*-like transposon in *P. aeruginosa* MRSN 12280. The isolate was non-susceptible to colistin by broth microdilution and genome analysis revealed no mutations known to confer colistin resistance. To the best of our knowledge this is the first report of *mcr* in colistin non-susceptible *P. aeruginosa*.

## Manuscript

*Pseudomonas aeruginosa* is a leading cause of infection among immunocompromised patients and those receiving treatment in Intensive Care Units (ICU) (1). Antimicrobial treatment of *P. aeruginosa* is challenging due to the intrinsic resistance of this species to many antibiotics and a proclivity to develop resistance to other antibiotics via point mutations in intrinsic genes (2, 3). Furthermore, *P. aeruginosa* can readily acquire transmissible antibiotic resistance (AbR) genes resulting in the emergence of successful multi- or extensively-drug resistant strains (4).The emergence of these resistant strains has resulted in a greater reliance on colistin (polymixin E) as a key antipseudomonal agent (5). Unfortunately, colistin resistance in *P. aeruginosa* has been extensively reported, primarily due to mutations in regulatory two-component systems (reviewed in (6)). However, colistin resistance mediated by the transferable colistin resistance gene *mcr* has not been described in this species to date. In this report we describe colistin non-susceptible *P. aeruginosa* MRSN 12280 carrying a chromosomal copy of *mcr-5*.*P. aeruginosa* MRSN 12280 was sequenced as part of a larger effort to sequence all *P. aeruginosa* isolates in the Multi-drug resistant organism Repository and Surveillance Network (MRSN) repository (n=2,440; manuscript in preparation). The isolate was cultured from a sacral wound of an elderly male patient treated in the USA in 2012. Minimum inhibitory concentrations (MICs) of colistin were determined using broth microdilution (BMD) with cation-adjusted Mueller Hinton broth (CA-MHB) according to the Clinical & Laboratory Standards Institute (CLSI) guidelines, and also with calcium-enhanced Mueller-Hinton (CE-MH) as recommended by Gwozdzinski and colleagues for *Enterobacteriaceae* carrying *mcr* (7). *Escherichia coli* MRSN 388734 carrying *mcr-1* (8) and *P. aeruginosa* ATCC 27298 were used as positive and negative controls, respectively. Colistin MICs were 4µg/ml (intermediate) and 8µg/ml (resistant) for *P. aeruginosa* MRSN 12280 in CA-MH and CE-MH medium, respectively (**Table 1**). Notably, the MIC of colistin in the control strains *E. coli* MRSN 388734 and *P. aeruginosa* ATCC 27298 also increased in CE-MH, but interpretations did not change (**Table 1**).

**Table 1.** Colistin susceptibility testing

| Strain | MIC of colistin in<sup>1</sup> |  |
| --- | --- | --- |
|  | CA-MH | CE-MH |
| *P. aeruginosa* MRSN 12280 | 4 | 8 |
| *E. coli* MRSN 388634 | 8 | 16 |
| *P. aeruginosa* ATCC 27298 | 0.25 | 1 |

Abbreviations used: MIC, minimum inhibitory concentration; CA-MH, cation-adjusted Mueller Hinton; CE-MH, Calcium-enhanced Mueller Hinton.
<sup>1</sup> MICs were performed in BMD according to CLSI guidelines. Results represent the average value from three independent tests.

Short and long-read whole genome sequencing (WGS) was performed on a NextSeq 550 (Illumina, San Diego, CA) and PacBio RS II (Pacific Biosciences, Menlo Park, CA), respectively, as previously described (9). *In silico* multi-locus sequence typing (MLST) assigned *P. aeruginosa* MRSN 12280 to a novel sequence type (ST) that is a single-loci variant of ST-2613. An analysis of the WGS data detected five AbR genes that are commonly found in *P. aeruginosa* (*aph(3′)-IIb*, *bla*_OXA-50_, *bla*_PAO_, *catB7*, and *fosA*) and the recently described colistin resistance gene, *mcr-5* (10). *Mcr-5* was first reported in 2017 in a cluster of colistin non-susceptible *Salmonella enterica* and shares a protein sequence identity of just 36.11% with Mcr-1 (10). Borowiak and colleagues reported that the gene was part of a Tn*3-*family transposon that was found primarily on small, multi-copy ColE-type plasmids. However, in one isolate (*S. enterica* 12-02546-2) the gene was present in a single copy on the chromosome and had a colistin MIC of 4mg/L (10). In *P. aeruginosa* MRSN 12280, a single copy of *mcr-5* was also present on the chromosome and was part of an 8,522bp Tn*3*-family transposon. This transposon was identical to the one described by Borowiak and colleagues except for the insertion of IS*5* into a gene encoding a putative major facilitator superfamily (MFS) directly downstream of *mcr-5* (**Figure 1**). The transposon was flanked by 38bp inverted repeats (IR) and generated a 5bp target site duplication (TCCAT) upon insertion.

**Figure 1.**
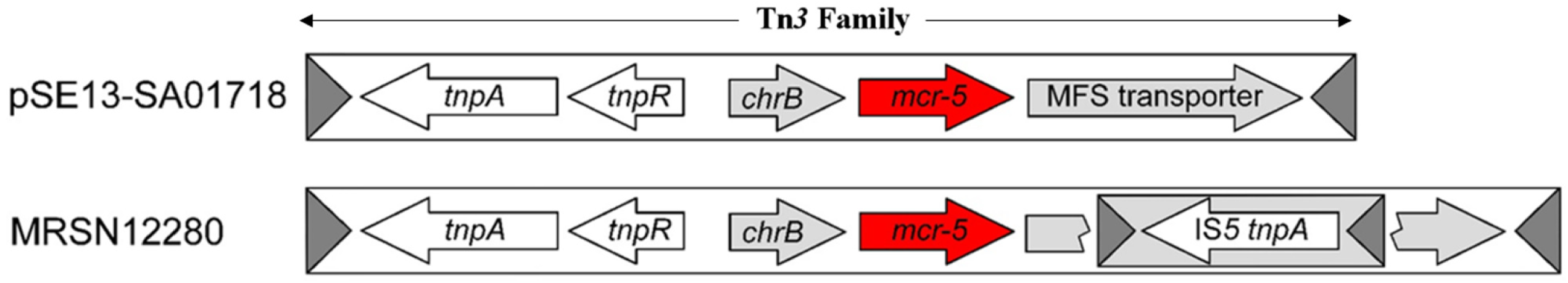
Alignment of Tn*3*-family transposons carrying *mcr-5*. Comparison of the Tn*3*-like transposon carrying *mcr-5* in plasmid pSE13-SA01718 from *Salmonella enterica* (10) with the Tn*3*-like transposon carrying *mcr-5* in *P. aeruginosa* MRSN 12280. Transposons are encased in a rectangle with the inverted repeats (IR) depicted as shaded, rotated triangles. Open arrows represent coding sequences (red arrows, *mcr-5*; white arrows, genes associated with DNA mobility; grey arrows, other genes) and indicate direction of transcription.

Colistin resistance in *P. aeruginosa* has primarily been attributed to mutations in up to five different two-component regulatory systems (PhoPQ, PmrAB, ParR/S, ColR/S, and CprR/S) (Reviewed in (6)) but to the best of our knowledge, Mcr-mediated colistin resistance has not been described in *P. aeruginosa* to date. As mutations in the two-component regulatory systems could potentially contribute to colistin resistance in *P. aeruginosa* MRSN 12280, we examined the amino acid sequences of PmrA, PmrB, PmrE, PhoQ, ParR, ParS, ColR, ColS, MigA, LpxC, and CprS for non-synonymous mutations (6, 11-13). When compared to *P. aeruginosa* PA01, *P. aeruginosa* MRSN 12280 had non-synonymous mutations in PhoQ (Y85F), PmrA (L71R), PmrB (S2P, A4T, G68S, Y345H, G362S), ParR (L153R, S170N), and ParS (H398R). However,all of the mutations in PhoQ, PmrA, and PmrB have previously been reported in colistin sensitive strains (12, 13). An analysis of 1,135 *parS* genes from the National Center for Biotechnology Information (NCBI) revealed that 1,117 sequences have an arginine at position 398, indicating that the ParS protein from *P. aeruginosa* PA01 is a poor representative of ParS in *P. aeruginosa*. Finally, a search of NCBI for the L153R and S170N mutations in ParR revealed that these mutations are present in a cluster of *P. aeruginosa* belonging to ST-235 from East Asia. *P. aeruginosa* VRFP04 from this cluster has been analyzed in detail and it is colistin sensitive (14). Though additional experiments are underway to confirm these findings, the data strongly suggests that colistin non-susceptibility in *P. aeruginosa* MRSN 12280 is due to Mcr-5.

We report the first identification of *mcr-5* in a colistin non-susceptible strain of *P. aeruginosa*. The gene was chromosomally encoded and embedded within a Tn*3-*family transposon that was related to the original transposon carrying *mcr-5* (10). Of note, while part of Mcr-5 and the transposon were identified in two *P. aeruginosa* assemblies using BLASTp (10), no other *P. aeruginosa* isolate in the MRSN repository contained the gene. This suggests that *mcr-5* is not widely distributed in this species but has the potential to disseminate via the Tn*3*-family transposon.

## Acknowledgements

This publication made use of the *Pseudomonas aeruginosa* MLST website (https://pubmlst.org/paeruginosa/) developed by Keith Jolley and sited at the University of Oxford (Jolley & Maiden 2010, BMC Bioinformatics, 11:595). The development of this site has been funded by the Wellcome Trust.

This study was funded by the U.S. Army Medical Command, the Global Emerging Infections Surveillance and Response System, and the Defence Medical Research and Development Program. Material has been reviewed by the Walter Reed Army Institute of Research. There is no objection to its presentation. The opinions or assertions contained herein are the private views of the authors and are not to be construed as official, or reflecting the views of the Department of the Army or the Department of Defence.

